# Designed allosteric protein logic

**DOI:** 10.1101/2022.06.03.494683

**Authors:** Tjaša Plaper, Estera Merljak, Tina Fink, Duško Lainšček, Tadej Satler, Vid Jazbec, Mojca Benčina, Roman Jerala

## Abstract

Regulation of the activity of proteins enables control of complex cellular processes. Allosteric regulation has been introduced individually into few natural proteins. Here, we present a generally applicable regulation of diverse proteins called INSRTR (inserted peptide structure regulator), based on inserting a short unstructured peptide into a solvent-accessible loop that retains protein function. Function of the target protein can be inactivated by the addition of a peptide that forms a rigid coiled-coil dimer. This platform enables the construction of ON/OFF protein switches, their regulation by small molecules, and Boolean logic functions with a rapid response in mammalian cells. INSRTR can be used to regulate a wide range of proteins, as demonstrated on ten members of protein families with diverse biological activities including enzymes, signaling mediators, DNA binders/transcriptional regulators, fluorescent protein, and antibodies regulating chimeric antigen receptor. INSRTR platform presents an extraordinary potential for regulating biological systems and applications.

**One sentence summary:** Authors have designed a widely applicable system to activate or inactivate function of diverse proteins or form Boolean logic gates based on formation of a coiled-coil dimer within protein loops and demonstrated its implementation on a range of 10 diverse proteins.

## INTRODUCTION

The regulation of protein function is a keystone of complex biological systems and is achieved through diverse mechanisms, including transcriptional regulation, multimolecular complex formation, posttranslational modifications, and allostery. So far, transcriptional regulation has dominated the design of engineered systems (Chavez et al., 2015; Church et al., 2014; Khalil and Collins, 2010; Verbič et al., 2020; Xie and Fussenegger, 2018) because regulation based on proteins is typically specific for each case and has been more difficult to engineer. Recently, several principles have been introduced for the engineering of protein function, such as the design of allosterically regulated proteins (Dagliyan et al., 2013, 2019; Ha and Loh, 2012; Karginov et al., 2010), the degradation of target proteins with degrons (CHOMP) (Gao et al., 2018), split protease and coiled-coil mediated reconstitution of split proteins (SPOC)(Fink et al., 2019), and the engineering of interaction domains for the alternative arrangement of the secondary structure elements (LOCKR) (Langan et al., 2019). Typically, these systems require extensive optimization for each individual protein, and their scope has been so far limited to a few selected protein types and functions. A widely applicable and robust platform for the regulation of protein function that could be introduced into diverse natural or engineered proteins would be highly desirable for research and applications that require control of in principle any selected protein.

A precisely defined tertiary structure is the prerequisite for the biological activity of proteins. However, high level of structural definition is often required only locally and encompasses a specific region of a protein, such as the catalytic site or the binding interface, while the rest of the protein structure serves as a scaffold to arrange the geometry of the functional site or support other functions. Proteins tolerate a high degree of sequence and structure variability particularly in solvent-exposed loops—and can often be genetically fused to other polypeptides while maintaining their function. Allostery, first named by Monod and Jacob in 1961 (Liu and Nussinov, 2016; Monod and Jacob, 1961), is a principle describing the regulation of proteins through a conformational change triggered by an effector binding at a site distal from the primary functional site. Although there are many allosterically regulated natural proteins, this process often involves an intricate relay of interactions and dynamics unique to each protein. Allosteric regulation has been introduced into proteins either by screening point mutations or through the insertion of a folded protein domain (Dagliyan et al., 2013, 2019; Ha and Loh, 2012; Ostermeier, 2005). We reasoned that it might be possible to introduce a general strategy of allosteric regulation to disrupt only the local conformation of diverse target proteins at a site crucial for their function, by altering the conformation of a solvent-exposed loop (**Figure 1A**). It is important that the platform could be extended, to enable construction of different types of switches and protein logic functions. Here, we present a strategy based on introduction of an intrinsically unstructured peptide at a permissible site of a host protein, hence enabling the protein to retain its function. This is generally feasible, because proteins often tolerate different loop lengths with divergent sequences. However, upon the binding of an appropriate binding partner of the inserted peptide, such as the complementary coiled-coil dimer peptide, the inserted peptide adopts an extended helical conformation, expanding the distance between the inserted termini in the loop; this is sufficient to locally disturb the conformation at the site crucial for protein function. We have implemented this concept as an inserted peptide structure regulator (INSRTR) platform with designed coiled-coil dimeric peptides as allosteric regulators. The wide applicability of this platform was demonstrated with a perfect success on ten different proteins comprising a wide range of protein folds and functions, including enzymes, signaling mediators, DNA-binding proteins as transcriptional regulators, and the single-chain variable fragment (scFv). While for most of these proteins allosteric regulation has not been reported before, we identified several permissible insertion sites, which governed the catalytic function or the binding interface of the host protein by adding a regulatory peptide provided through different means into mammalian cells. Typically, several permissible insertion sites were identified with a high success rate within each protein, which retained the function but could be controlled in mammalian cells by adding a regulatory peptide provided through different means. The scope of this platform was further extended by a genetic fusion with an intramolecular inhibitory peptide to invert the regulation from an OFF to an ON switch and to trigger regulation by small molecules acting via a chemically regulated split protease; this further enabled the construction of Boolean logic functions in mammalian cells. The wide diversity of applicable targets and the ease of construction demonstrate that this principle has remarkable potential for engineering proteins to regulate natural or engineered biological systems.

**Figure 1.**
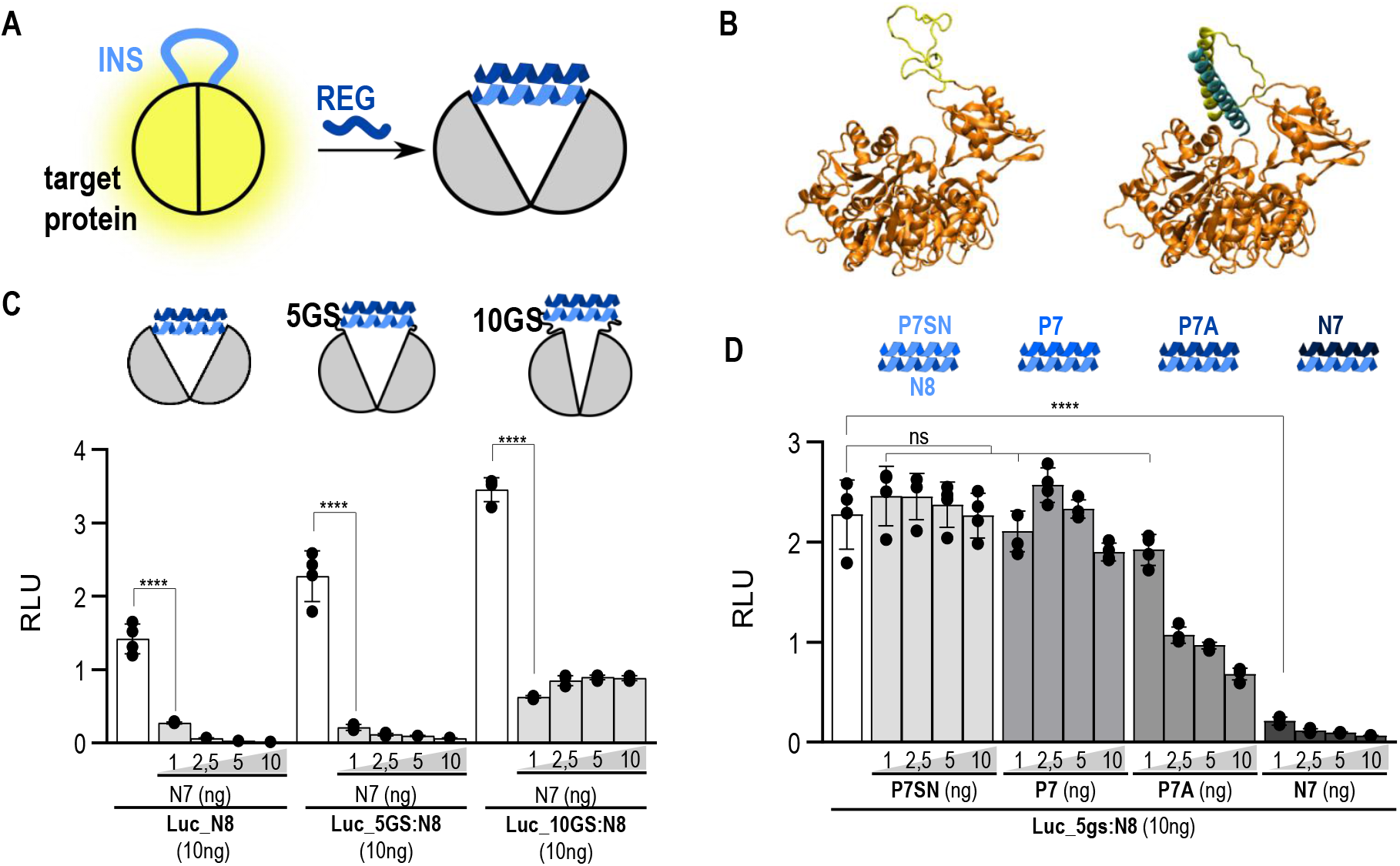
The principle of Inserted peptide structure allosteric regulation (INSRTR) of target protein function. **(A)** Scheme of the INRSTR principle, unstructured inserted peptide maintains structure and function, coiled-coil formation triggered by the regulatory peptide; INS-inserted peptide, REG-regulatory peptide. **(B)** Molecular model of INSRTR firefly luciferase that maintains its structure with insertion of the unstructured CC-forming peptide (inserted peptide) into the loop. The addition of a regulatory peptide leads to coiled-coil formation and thus structural changes in firefly luciferase leading to its inactivation; fLuc-firefly luciferase. **(C)** Peptide N8 was genetically fused into firefly luciferase with different lengths of connecting linkers on both sides of the peptide sequence. Linker regions allow for additional flexibility around peptide which effects maintained firefly luciferase activity as well as the level of inhibition with the addition of peptide pair. **(D)** The level of inhibition is dependent on CC pair affinity. Values in B and C are the mean of four biological replicates ± (s.d.) and representative of three independent experiments on plasmid transfected HEK293T cells. Significance was tested with one-way ANOVA with Dunnett’s multiple comparisons test between INSRTR variant and addition of a REG peptide; p values 0.1234 (ns), 0.0332 (*), 0.0021 (**), 0.0002 (***), <0.0001 (****) (significance, confidence intervals, degrees of freedom, F and P values are listed in Table S2).

## RESULTS

### The principle of inserted peptide structure allosteric regulation (INSRTR)

Because the main impact of introducing protein allosteric regulation lies in biological systems, the concept was explored in mammalian cells. The insertion sites were selected based on the following considerations: a) position within the solvent exposed loop, b) no residues are directly involved in protein function, c) are preferentially variable in length and sequence in homologues, c) loop is connected to the secondary structure element that is coupled through a network such as the super secondary structure to the functional site, and e) position is separated from the functional site by 1-4 nm.

The principle initially tested on the firefly luciferase, which is a suitable reporter to establish the principles that could be applied to other proteins. Based on the 3D structure we identified candidate insertion sites. Position 490 was selected, separated 20 Å from the proposed active site (Conti et al., 1996). For allosteric regulators, designed coiled-coil (CC) dimer peptide pairs—orthogonal to natural leucine zippers—and affinity tunable in the nano-to micromolar range were used (Drobnak et al., 2017a; Lebar et al., 2020; Plaper et al., 2021). A CC dimer-forming peptide that is unstructured in the absence of the binding partner was inserted at selected position (**Figure 1B, left**). In the presence of the regulatory peptide, a CC dimer should form (**Figure 1B, right**), with the inserted peptide segment adopting an extended helical conformation with a distance between its termini of ∼4 nm. This disrupts the local structure of the protein, which could affect the geometry of the active site. Indeed, co-expression of the regulatory peptide resulted in the strong inhibition of luciferase activity (**Figure 1C**). Molecular dynamics simulation of the luciferase model (**Figure 1B**) suggested that the formation of a CC dimer in the loop affected the conformation of the loop, while the rest of the host protein, including the geometry and dynamics of the active site residues, remained essentially unperturbed (Figure not shown). Therefore, inhibition is likely a result of a subtle perturbation of the structure or/and dynamics of the active site.

To optimize the ratio between an active and inactive state in the absence or presence of a regulatory peptide, respectively, the length of the linker peptide between the CC-forming insert and insertion site of the protein was varied. This revealed an optimal response for a five-residue linker peptide with up to a 30-fold repression of activity in the presence of the regulatory peptide (**Figure 1C**). Although the absence of a flexible linker suppressed the activity of an active state, longer linker, on the other hand, caused only partial abrogation of activity since long flexible linker decouples the conformational transition of the inserted peptide from the rest of the host protein. CC pairs can be designed in a wide range of stabilities (Drobnak et al., 2017b) and have been shown to be orthogonal to the endogenous CCs (Lebar et al.); indeed, a comparison of four CC pairs revealed that the degree of inhibition was proportional to their affinities (Drobnak et al., 2017b; Plaper et al., 2021) (**Figure 1D**). Hence, the response of the allosteric switch could be tuned using a toolbox of CC dimer pairs with different affinities (Drobnak et al., 2017a).

### Designed inverted INSRTR (ON switch) can be regulated by various trigger signals

The strategy presented so far allows inhibition of protein activity through the addition of a regulatory peptide. Often, a constitutively inactive protein is desired that could be activated by a selected trigger. To design an inverted ON variant of INSRTR, an inhibitory peptide with a moderate affinity to the inserted peptide was genetically fused to the C-terminus of the host protein (**Figure 2A**). This enabled the intramolecular binding of inhibitory peptide to the inserted peptide and a constitutively inactive state of a target protein. A regulatory peptide with a higher affinity for the inhibitory peptide could compete for binding and release the inserted peptide from an intramolecular dimer, hence restoring the activity of the host protein, which represents an ON switch (**Figure 2A**). This concept was demonstrated on a luciferase with inserted P7 peptide and a C-terminal fusion of moderate-affinity N8 inhibitory peptide (Plaper et al., 2021). The formation of an intramolecular P7-N8 heterodimer resulted in an inhibited luciferase. The addition of a regulatory N7 peptide with high affinity for the N8 inhibitory peptide (Plaper et al., 2021) recovered luciferase activity (**Figure 2B**). Several modes of regulation of allosteric ON switch in mammalian cells were implemented, including transcriptional regulation of peptide expression (**Figure 2B**), external delivery of the peptide to cells (**Figure 3A**), or proteolytic cleavage of the linker between the host protein and the inhibitory peptide (**Figure 3B**). For the latter, tobacco etch virus protease (TEVp) was used, which could be activated by rapamycin-inducible heterodimerization between FKBP and FRB fused to the split protease (Fink et al., 2019). The P4 peptide—which has a different electrostatic and hydrophobic motif when compared with N8 (Plaper et al., 2021) and that, therefore, does not bind to N8—had no effect on luciferase activity (**Figure 3A**), supporting the orthogonality of the designed CC pairs. Since the INSRTR relies on protein–protein interactions, it was expected that the addition of a regulatory peptide could trigger a fast response in mammalian cells. Indeed, luciferase activity was observed within 5 minutes of exposing mammalian cells to rapamycin (**Figure 3C**). In this way, a small molecule chemical trigger can be used to regulate the function of selected allosterically regulated proteins in cells.

**Figure 2.**
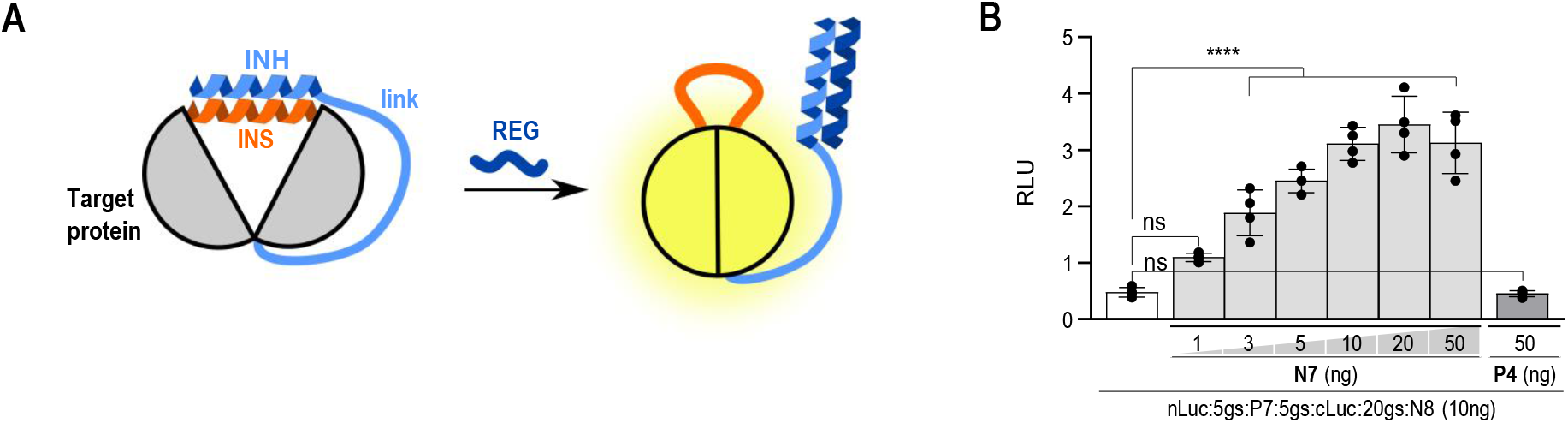
Design and characterization of inverted INSRTR and formation of an ON switch. **(A)** Schematic presentation of inverted INSRTR (ON switch) design. The ON switch was designed by genetic fusion of a low-affinity CC peptide (inhibitory peptide) to the C-terminus of the target protein and into the protein loop (inserted peptide). Because of the high local concentration, the inhibitory and inserted peptides form an intramolecular CC dimer, resulting in an inactivated target protein. The addition of the regulatory peptide, with high affinity for CC formation with an inhibitory peptide, results in an unstructured inserted peptide, thus regaining target protein function. Different N7 peptide delivery systems were evaluated on the ON-INSRTR system. **(B)** Activity of ON-INSRTR firefly luciferase is upregulated with the co-expression of REG peptide N7, forming the designed CC pair. Presence of orthogonal CC-forming peptide P4 does not affect firefly luciferase activity. The values in B are the mean of four biological replicates ± (s.d.) and representative of at least two independent experiments on transiently transfected HEK293T cells. Significance was tested with one-way ANOVA with Dunnett’s multiple comparisons test between INSRTR variant and addition of a REG peptide; p values 0.1234 (ns), 0.0332 (*), 0.0021 (**), 0.0002 (***), <0.0001 (****) (significance, confidence intervals, degrees of freedom, F, and P values are listed in **Table S2**).

**Figure 3.**
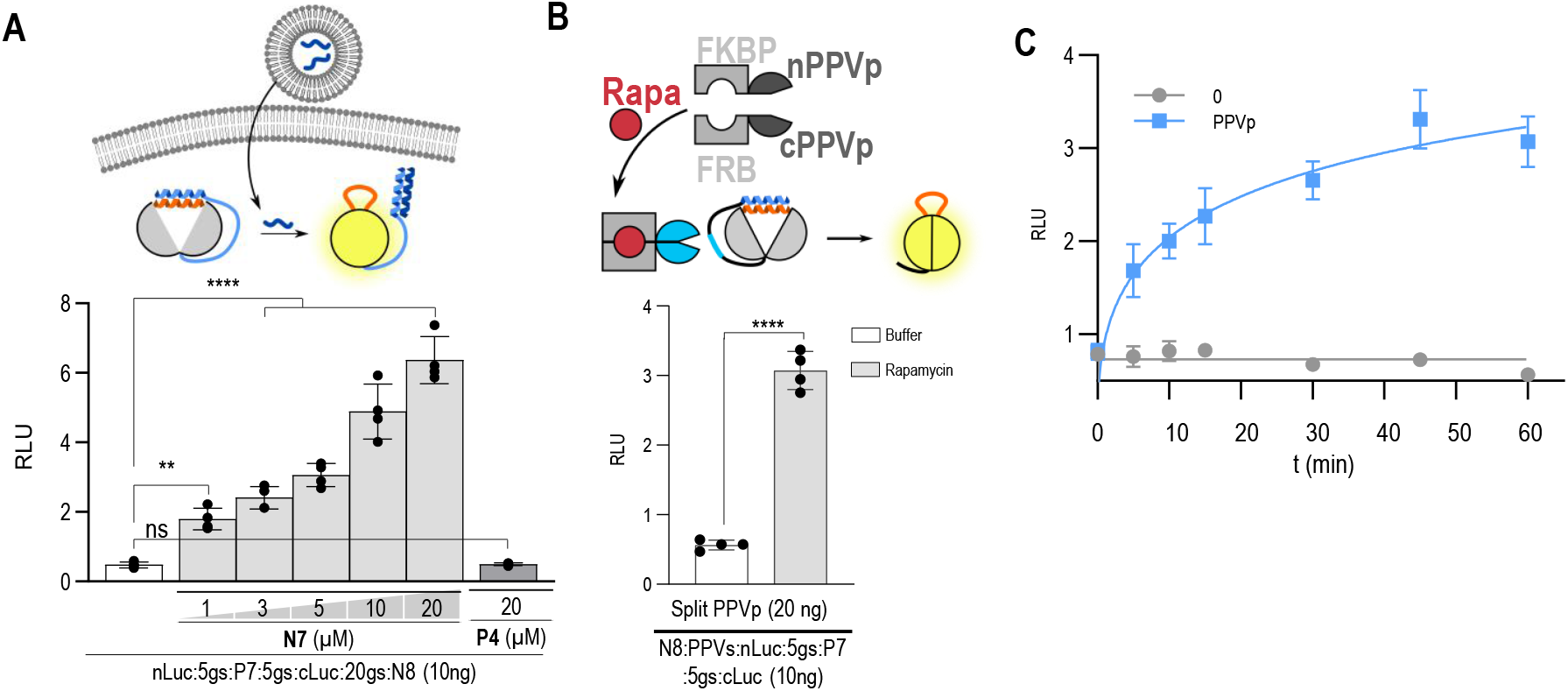
Different modes of providing a trigger of INSRTR and kinetics in mammalian cells. **(A-B)** Two other N7 peptide delivery systems were evaluated in addition to transcriptional regulation on the ON-INSRTR system by **(A) an** externally added peptide, where an orthogonal CC-forming peptide P4 does not affect the firefly luciferase activity. **(B)** Chemically activated protease regulation of ON-INSRTR switch mediated by cleavage of the linker peptide between INH and the target protein. **(C)** Regulation of rapamycin-mediated reconstitution of TEV protease results in the fast kinetics of ON-INSRTR system activation. The values in (A-C) are the mean of four biological replicates ± (s.d.) and representative of at least two independent experiments on transiently transfected HEK293T cells. Significance was tested with one-way ANOVA with Dunnett’s multiple comparisons test between INSRTR variant and addition of a REG peptide; p values 0.1234 (ns), 0.0332 (*), 0.0021 (**), 0.0002 (***), <0.0001 (****) (significance, confidence intervals, degrees of freedom, F, and P values are listed in **Table S2**). INS-inserted peptide, INH-inhibitory peptide, REG-regulatory peptide, link-linker peptide, Rapa-rapamycin. See also **Figure 2A** for schematic presentation of activation and **Figure 2B** for comparison to co-expression of REG peptides.

### Construction of Boolean protein logic gates based on intramolecular interacting segments

The concept of intra- and intermolecular interactions with regulatory CC segments and proteolysis can be extended to the construction of a full set of Boolean logic functions, similar to SPOC logic (Fink et al., 2019) (**Figure 4A-B, Figure S1**). The input signals are provided through orthogonal proteases, whose activity can be regulated by different small molecules. Other types of input signals could be proteases specific to certain biological processes, for example, caspases or viral proteases. We designed constructs with CC peptides fused to the C or N termini of firefly luciferase (**Figure 4C**). By implementing designable CC peptides with appropriate affinities and cleavage sites for selected proteases, we were able to implement all combinations of two signal inputs (**Figure 4C**). A particular advantage of this strategy is that logic functions were genetically encoded within a single polypeptide chain using two orthogonal proteases activity as the input signals, which could be regulated by small molecules. Intracellular fusion with CC-forming peptides, in contrast to multiple chain designs, also ensured a faster response (**Figure 4D**).

**Figure 4.**
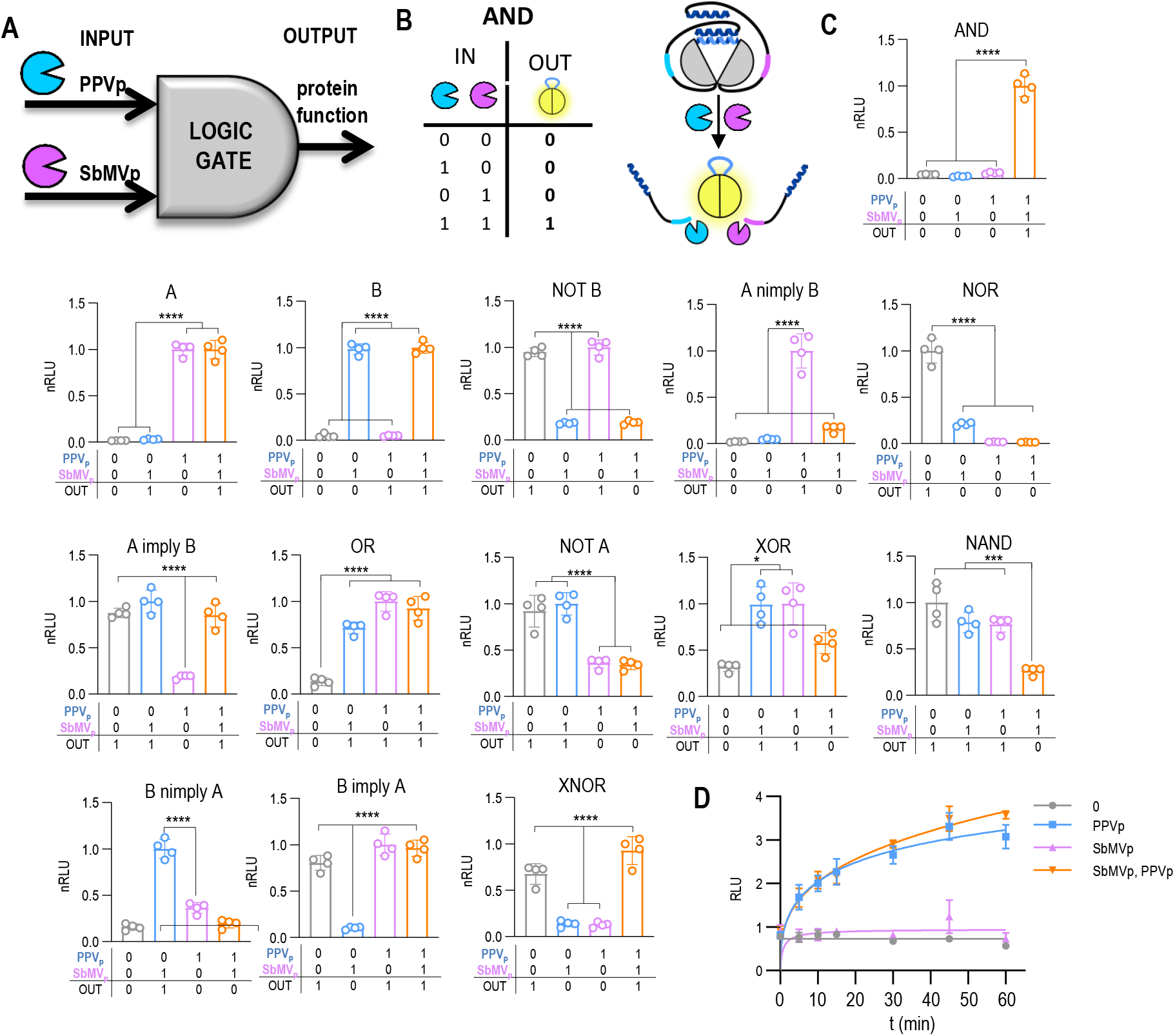
Construction of Boolean protein logic gates based on intramolecular interacting segments. **(A) Scheme of the** implementation of INSRTR logic gates with protease inputs and allosterically activated protein function. **(B)** Schematic presentation of an AND logic gate. **(C)** Experimental testing of all nontrivial two-input Boolean logic functions by INSRTR in HEK293T cells, as regulated by a combination of PPV and SbMV protease inputs. Schemes of the constructs are provided in the **Figure S1**. **(D)** Kinetics of a B function response by chemical regulation of the activity of protease inputs. The values in C and D are the mean of four cell cultures ± (s.d.) and are representative of at least two independent experiments. Significance was tested with one-way ANOVA with Tukey’s comparison between indicated ON and OFF states. p values 0.1234 (ns), 0.0332 (*), 0.0021 (**), 0.0002 (***), <0.0001 (****) (significance, confidence intervals, degrees of freedom, F, and P values are listed in **Table S2**). **See also Figure S1**.

### INSRTR regulation of diverse protein types and functions in an OFF or ON format

The most important impact of INSRTR as an allosteric regulator was found to be its ability to control the function of a range of diverse proteins, which opens exciting possibilities to modulate biological systems, such as e.g. to temporally (in)activate DNA cleavage or suppress excessive activation of T cells. INSRTR-permissible sites are likely to be available in many different proteins. To showcase the wide applicability of INSRTR, we demonstrated it on ten different proteins comprising different superfamilies with diverse biological functions and quaternary structures, including enzymes (protease, β-galactosidase, kinases), signaling proteins (signaling adaptor, kinases), transcriptional regulators based on DNA-binding domains (transcription activation-like effector (TALE) and Cas9), fluorescent proteins, and antibody. Among the enzymes, in addition to the luciferase, INSRTR was successfully implemented into TEVp, β-galactosidase, Lck, and IRAK1 protein kinases (**Figure 5A-E**). For each protein, at least one—but often up to three—sites were found to be amenable to the INSRTR platform (**Figure S2-S4**). Activity of each protein was evaluated by its specific biological assay. Upon co-expression with regulatory peptide we achieved up to 90% inhibition. For all four enzymes, we successfully used regulatory peptides to downregulate the activity (OFF) and, in most cases, an autoinhibited version that could be turned ON by the addition of a regulatory peptide. An important group of proteins are mediators that regulate signal transduction in the immune response, here either through kinase activity, such as Lck or IRAK1 kinase, or as signaling adaptors recruiting other signaling components, such as MyD88. In the case of the IRAK1 kinase, a peptide was introduced into the catalytic domain of the kinase. The addition of a regulatory peptide completely inhibited the signaling function of the kinase. (**Figure 5E**). In the case of signaling adaptors that function through protein– protein interactions, sensitivity to the allosterically induced strain in the structure might have a less potent effect than on catalytic activity, where the active site architecture must be defined with high precision. Nevertheless, the function of a key innate immune signaling mediator, MyD88, which interacts with Toll-like receptors and IL-1 receptor via its Toll-interleukin receptor (TIR) domain and recruits downstream kinases by a death domain (DD) (Avbelj et al., 2011), was successfully regulated by a peptide through modulation of the binding interface (**Figure 5F**). Three permissible sites within the TIR domain and one within a DD of MyD88 were identified (**Figure S3A-D**). An important group of proteins are transcriptional regulators that bind to a specific DNA sequence based on recognition by the protein (TALE proteins (Mak et al., 2012), **Figure 5G**) or through an RNA-mediated recognition through a Cas9/gRNA complex (Jinek et al., 2014) (**Figure 5H, Figure S4A-D**). In both cases, we successfully disrupted the DNA recognition interface through the binding of a regulatory peptide to the inserted peptide, introducing a new control for those types of transcriptional regulators. Similar to other proteins, we were able to prepare either an OFF or ON TALE or dCas9-mediated transcriptional activators (**Figure 5G-J**). Next, we demonstrated that GFP fluorescence could be regulated by insertion at position 143 (**Figure 5K, Figure S5**), which might therefore also serve as a peptide sensor. Finally, antibodies are physiologically important players in adaptive immunity but also as reagents targeting many proteins and other cellular components or as therapeutics. The ability to regulate the binding of antibodies to their targets creates opportunities for research and applications. The variable domains of a single chain antibody (scFv) are used in chimeric antigen receptors (CAR) for the activation of T cells and elimination of target cancer cells (Srivastava and Riddell, 2015). Excessive activation of CAR T cells may lead to a cytokine storm and even a lethal outcome (Srivastava and Riddell, 2015). Therefore, it is desirable to temporally and rapidly desensitize CAR T cells rather than to eliminate them via a kill switch. Inserting a peptide at position 195 of the recognition domain scFv did retained binding and activation of T cells in the presence of CD19-expressing target cells, while the addition of a regulatory peptide strongly suppressed the response of CAR T cells. This principle could therefore be implemented to prevent adverse therapeutic effects (**Figure 5L, Figure S4E-H**).

**Figure 5.**
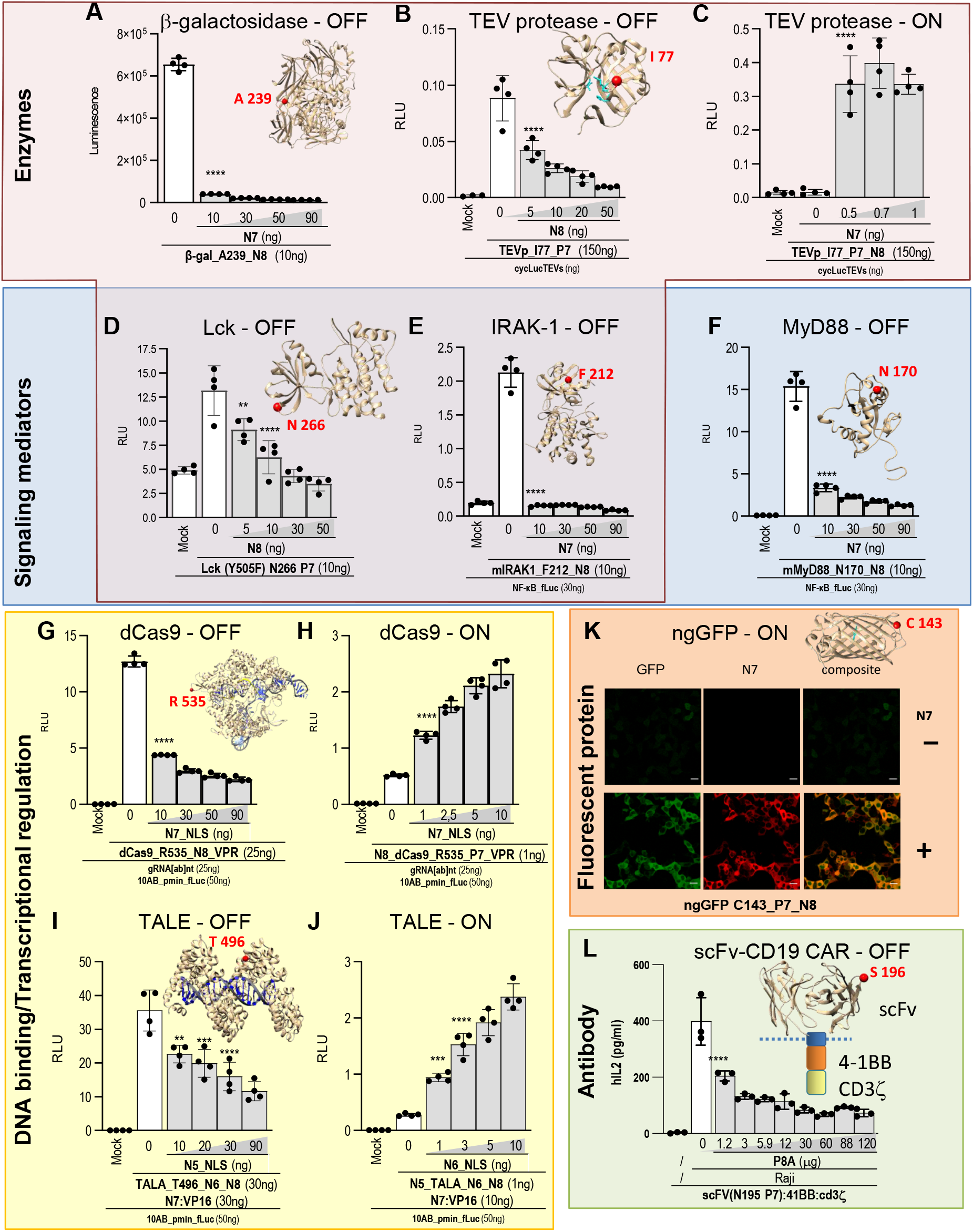
INSRTR regulation of diverse protein types and functions in an OFF or ON format. **(A-L)** Evaluation of INSRTR regulation on representative proteins from various protein groups. Enzymes: **(A)** β-galactosidase, **(B-C)** TEV protease, **(D)** Lck kinase; Signaling mediators **(E)** IRAK1, **(F)** MyD88; DNA binding-based transcriptional regulators **(G-H)** dCas9, **(I-J)** TALE; Fluorescent proteins **(K)** GFP and CAR-T receptor **(L)** scFv-CD19. Inserts are schematic presentations of allosterically regulated protein 3D structures with highlighted site of peptide insertion (red sphere). Mock represents transfection with reporter plasmids only. Values in (A-K) are the mean of four biological replicates ± (s.d.) and representative of three independent experiments on transiently transfected HEK293T cells. Significance was tested with one-way ANOVA with Dunnett’s multiple comparisons test between INSRTR variant and addition of a REG peptide; p values 0.1234 (ns), 0.0332 (*), 0.0021 (**), 0.0002 (***), <0.0001 (****) (significance, confidence intervals, degrees of freedom, F and P values are listed in **Table S2**). **See also Figure S2-S5**.

## DISCUSSION

INSRTR platform represents a generally applicable principle for introduction of regulation into the selected proteins and their coupling to chemical inputs. Insertion positions were selected based on several rational considerations, similar as before (Dagliyan et al., 2019), however the success rate was comfortably above 30% for most proteins. We successfully implemented INSRTR into each of the ten proteins, demonstrating the robustness of this platform. While, activity of target proteins varied across selected position of insertion, we were able to prepare for all investigated proteins at least one variant of with high activity and large fold inhibitions upon regulatory peptide expression. Stronger affinity between the regulatory peptides correlated with the more potent response. Coupling of INSRTR to chemically regulated proteases facilitates regulation of diverse proteins by small molecules. INSRTR enables rapid response inside mammalian cells via the control of protein function and construction of interaction networks and logic operations. This platform requires a small genetic footprint of the insert and is expected to have low immunogenicity because of the designability of CC peptides (Drobnak et al., 2017b; Fletcher et al., 2012; Reinke et al., 2010). Implementing INSRTR requires minimal engineering while providing a high success rate; in many cases, INSRTR permits several different insertion sites that could be important to bypass interference through any other interaction interfaces. The toolbox of designed CC pairs has been shown to be tunable and orthogonal in mammalian cells (Lebar et al., 2020). The regulation of INSRTR can be implemented through transcriptional activation, small molecules, or the addition of an external peptide. The ON or OFF adaptation of a target protein offer the possibility to engineer complex regulatory circuits with a fast response.

Allosteric regulation has been introduced before into proteins through the insertion of folded domains (Ha and Loh, 2012), with proteins such as FKBP, DHFR and ubiquitin (Karginov et al., 2010; Oakes et al., 2016; Radley et al., 2003) or through light-inducible domains (Strickland et al., 2008). However, the effect of domain insertion has been largely unpredictable (Stein and Alexandrov, 2015). The insertion sites of an FKBP domain have been investigated for Cas9 by an extensive library screen (Oakes et al., 2016), while our design identified previously unidentified sites permissible to allosteric regulation. INSRTR can be seen as a protein analogy of a transistor that amplifies the binding signal at a distal site through allosteric coupling to the protein function, with a range of the response up to 30 fold.

Importantly, diverse types of proteins and functions can be regulated through INSRTR switches. It has been previously shown that CC interactions can be regulated by competitive binding, antibodies, metal ions (Aupič et al., 2018), phosphorylation, and proteolysis (Fink et al., 2019); therefore, several signals and processes could be plugged into the INSRTR platform. The wide range of protein structures and functions admissible for regulation by INSRTR opens up a broad range of possibilities for regulating biological processes.

### Limitations of the study

Although we have demonstrated INSRTR only in mammalian cells, the basic biochemical principles should make it feasible in other biological or *in vitro* systems. It is likely that some hub proteins that interact with many partners along several sites [33] or intrinsically disordered proteins might be difficult to engineer without interference of some interactions, nevertheless engineering of MyD88 as a central mediator of Toll-like receptor signaling involved in the multiprotein complexes s was successful. The study was performed in mammalian cells and it will be interesting to confirm the mechanism of action by *in vitro* structural studies in the future. Nevertheless, in most cases several insertion positions in different loops were successful, which confirms that the inhibition is not based on steric hindrance, which is further supported by the fact that longer flexible linkers abolished regulation and that we could engineer also enzymes with small substrates. Additionally, fast activation in mammalian cells also supports the proposed mechanism. It is likely that some hub proteins that interact with many partners along several sites (Perica et al., 2021) or intrinsically disordered proteins might be difficult to engineer, nevertheless engineering of MyD88 as a central mediator of Toll-like receptor signaling involved in the multiprotein signaling complex was successful.

## Acknowledgments

The project was funded by the Slovenian Research Agency (program P4-0176, projects J1-9173, Z4-2657, N4-0800) and European Research Council (ERC AdG MaCChines 787115). We thank Ajasja Ljubetič and Kristina Djinović Carugo for fruitful discussions and comments.

## Author Information

These authors contributed equally: T.P., E.M., T.F.

## Author contribution

Conceptualization, R.J.; Formal analysis, T.P., E.M., and T.F.; Funding acquisition, R.J.; Investigation, T.P., E.M., T.F., and D.L., Methodology, T.S.; Supervision, R.J.; Validation, V.J., M.B.; Visualization, T.P., E.M., and T.F.; Writing – original draft, R.J.; Writing – review & editing, R.J., T.P., E.M., and T.F.

