## Supplemental file for "Designed allosteric protein logic"

**This PDF file includes:**

Fig S1-S5

Materials and Methods

### SUPPLEMENTAL FIGURE TITLES AND LEGENDS

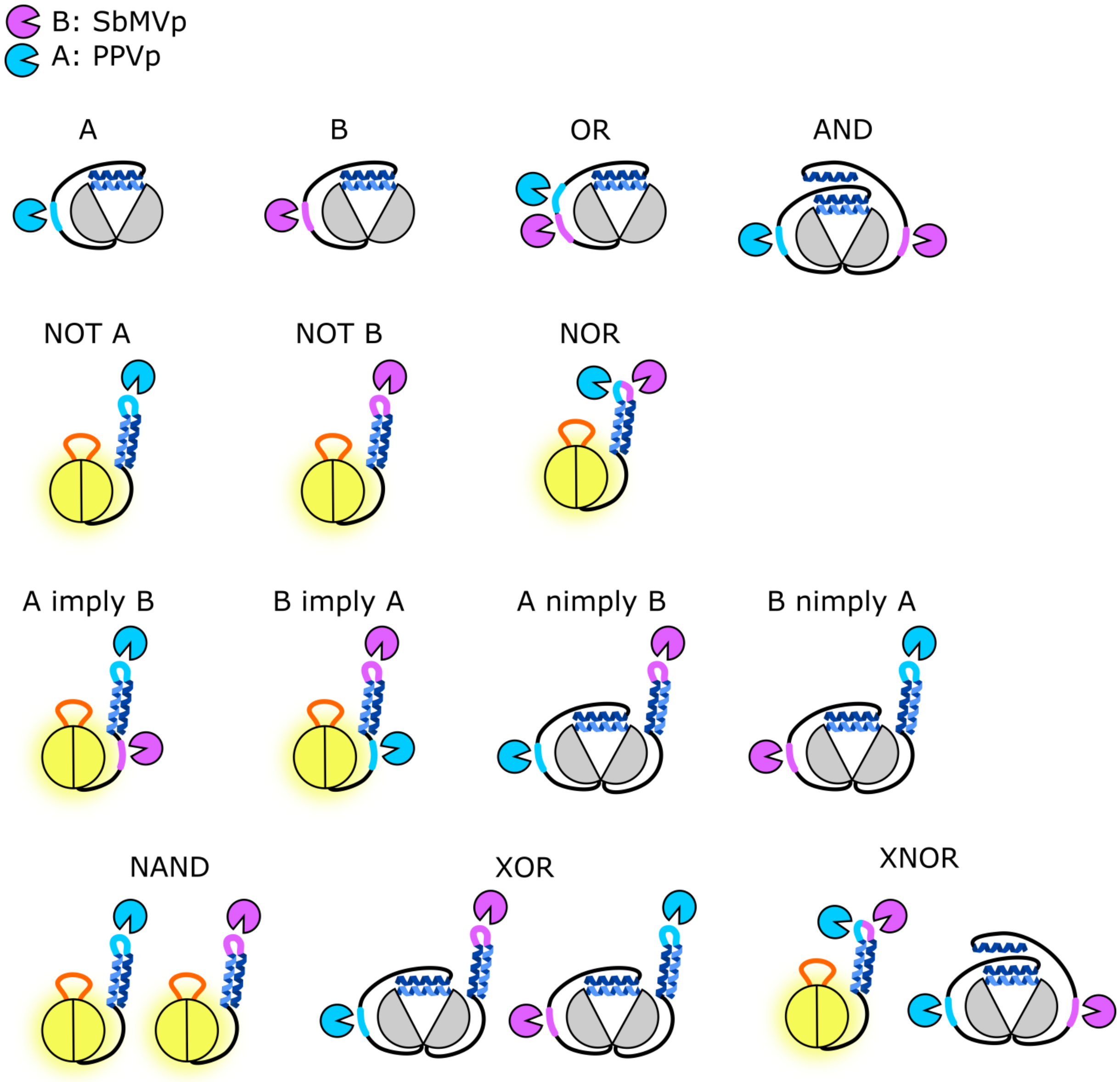


**Figure S1. Schematic representation of INSRTR firefly luciferase used for construction of all Boolean gates based on cleavage of intramolecular inhibitory segments. Related to Figure 4.**

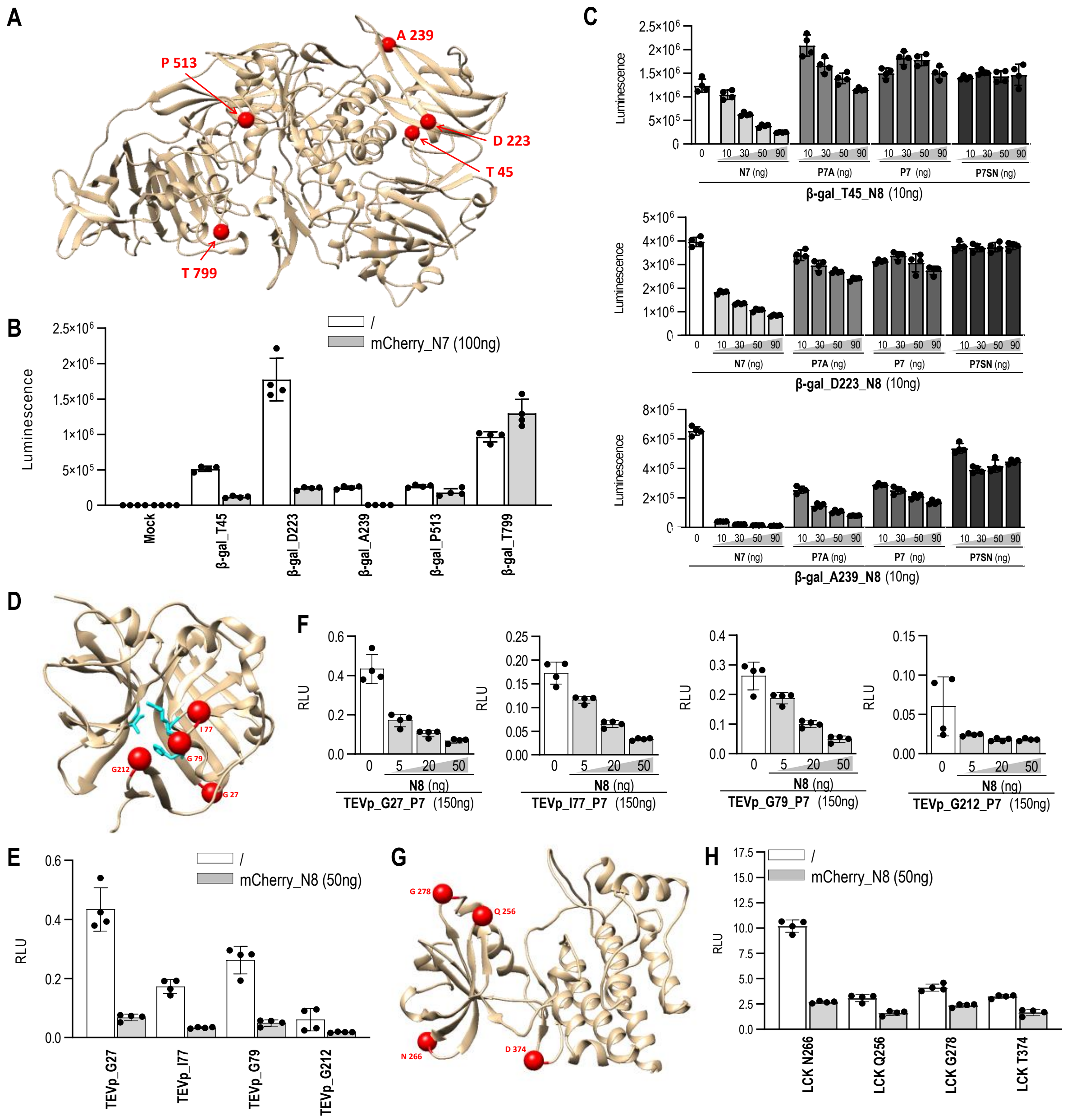


**Figure S2. Characterization of β-galactosidase, TEVp and Lck INSRTR variants. Related to Figure 5.**

**(A)** Structure of β-galactosidase with highlighted sites of N8 peptide insertion (red).

**(B)** Activity of designed β-galactosidase INSRTR versions in the absence or presence of 100 ng of plasmid encoding for mCherry_N7.

**(C)** Titration of plasmids encoding different affinity variants of peptides: N7, N7A, P7 and P7SN on three β-galactosidase INSRTR variants.

**(D)** Structure of TEVp with highlighted sites of N8 peptide (red).

**(E)** Activity of designed TEVp INSRTR versions in the absence or presence of 50 ng of plasmid encoding for mCherry_N7.

**(F)** Titration of plasmids encoding for N7 peptide on four TEVp INSRTR variants.

**(G)** Structure of Lck with highlighted sites of N7 peptide insertion (red).

**(H)** Activity of designed Lck INSRTR versions in the absence or presence of 50 ng of plasmid encoding for mCherry_N8.

Values in B, C, E, F and H are the mean of four biological replicates ± (s.d.) and representative of three independent experiments on transiently transfected HEK293T cells.

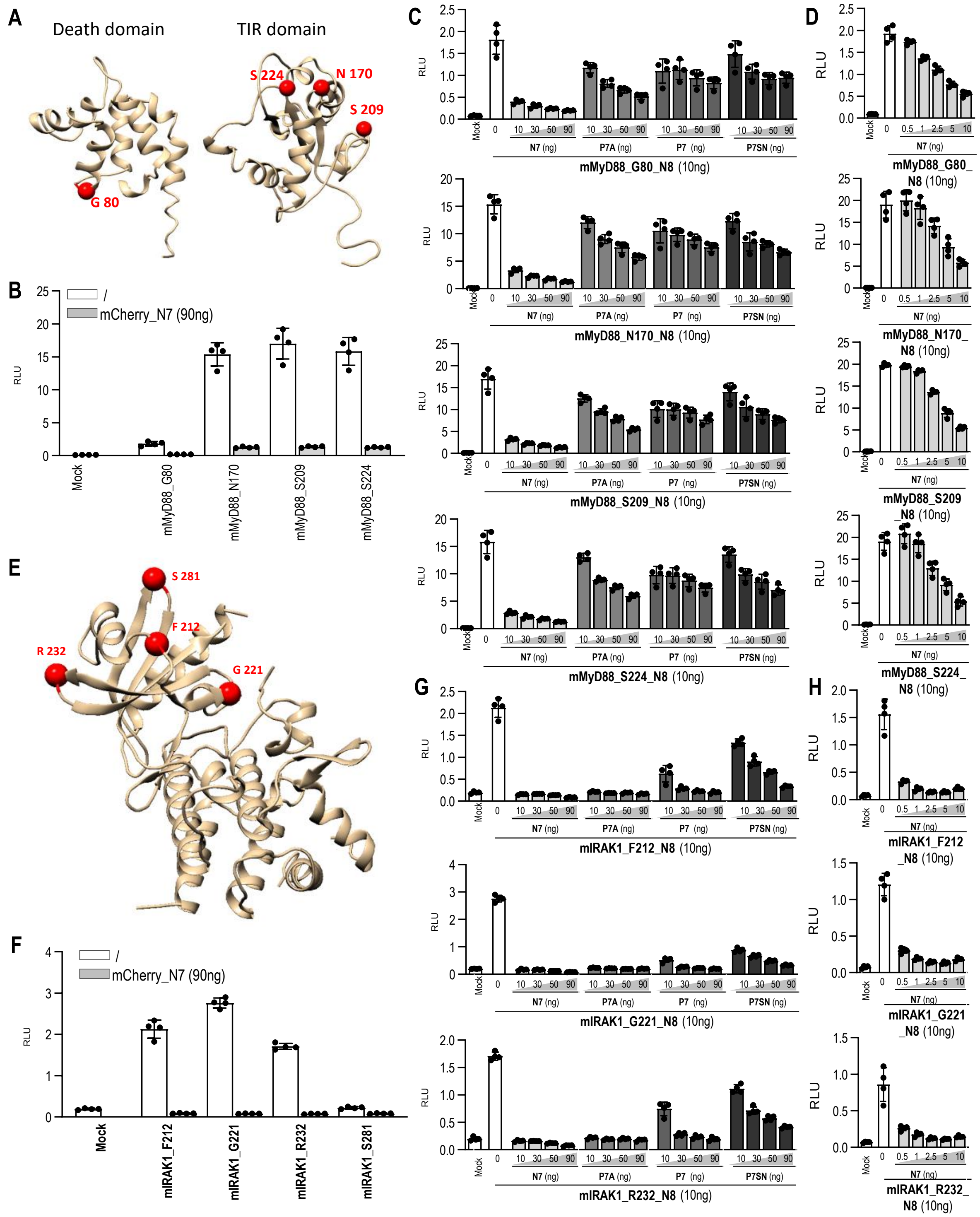

**Figure S3. Characterization of mMyD88 and mIRAK1 INSRTR variants. Related to Figure 5.**

**(A)** Structure of hMyD88 death domain and TIR domain with highlighted sites of N8 peptide insertion (red). G80, S209 and S224 are conserved among human and mouse MyD88. The site S170 in hMyD88 is represented as N170 in mMyD88.

**(B)** Activity of designed mMyD88 INSRTR versions in the absence or presence of 90 ng of plasmid encoding for mCherry_N7.

**(C)** Titration of plasmids encoding different affinity variants of peptides: N7, N7A, P7 and P7SN on mMyD88 INSRTR variants.

**(D)** Titration with lower plasmids amounts encoding for N7 peptide on mMyD88 INSRTR variants.
**(E)** Structure of hIRAK1 with highlighted sites of N8 peptide insertion (red). F212, G221 and R232 are conserved among human and mouse IRAK1. The site N281 in hIRAK1 is represented as S281 in mIRAK1.
**(F)** Activity of designed mIRAK1 INSRTR versions in the absence or presence of 90 ng of plasmid encoding for mCherry_N7.
**(G)** Titration of plasmids encoding different affinity variants of peptides: N7, N7A, P7 and P7SN on three mIRAK1 INSRTR variants.
**(H)** Titration with lower plasmids amounts encoding for N7 peptide on three mIRAK1 INSRTR variants.
Values in B-D, and F-H are the mean of four biological replicates ± (s.d.) and representative of three independent experiments on transiently transfected HEK293T cells.

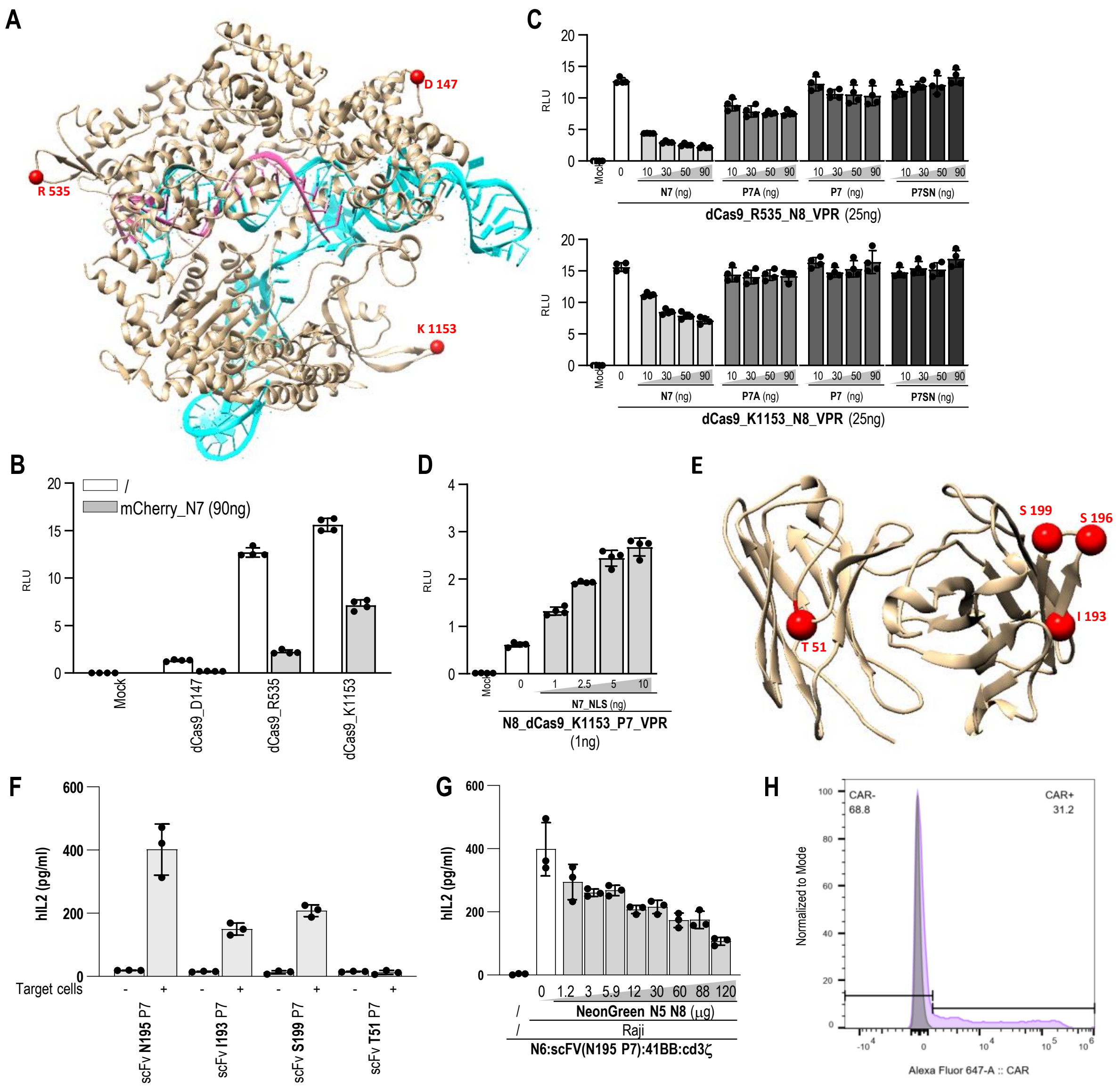

**Figure S4. Characterization of dCas9 and scFvCD19 INSRTR variants. Related to Figure 5.**

**(A)** Structure of dCas9 with highlighted sites of N8 peptide insertion (red). DNA is shown in cyan; gRNA in magenta.

**(B)** Activity of designed dCas9 INSRTR versions in the absence or presence of 90 ng of plasmid encoding for mCherry_N7.

**(C)** Titration of plasmids encoding different affinity variants of peptides: N7, N7A, P7 and P7SN on two dCas9 INSRTR variants.

**(D)** Induction of dCas9_K1153 INSRTR with N7 peptide.

**(E)** Structure of scFvCD19 with highlighted sites tested for P7 peptide insertion (red).

**(F)** Activity of designed INSRTR variants of scFv CD19 CAR-T.

**(G)** Inhibition of Jurkat T cell activation by the addition of B cells (Raji) by the addition of a coiled-coil forming peptide. Insertion into scFv at position 196 was responsive to the addition of a protein containing N8 coiled-coil forming peptide.

**(H)** Flow cytometry showing surface expression of INSRTR CAR-T. Cells were stained with 9B11 anti Myc-tag AF647 antibody.

Values in B-D and F-G are the mean of four biological replicates ± (s.d.) and representative of three independent experiments on transiently transfected HEK293T cells.

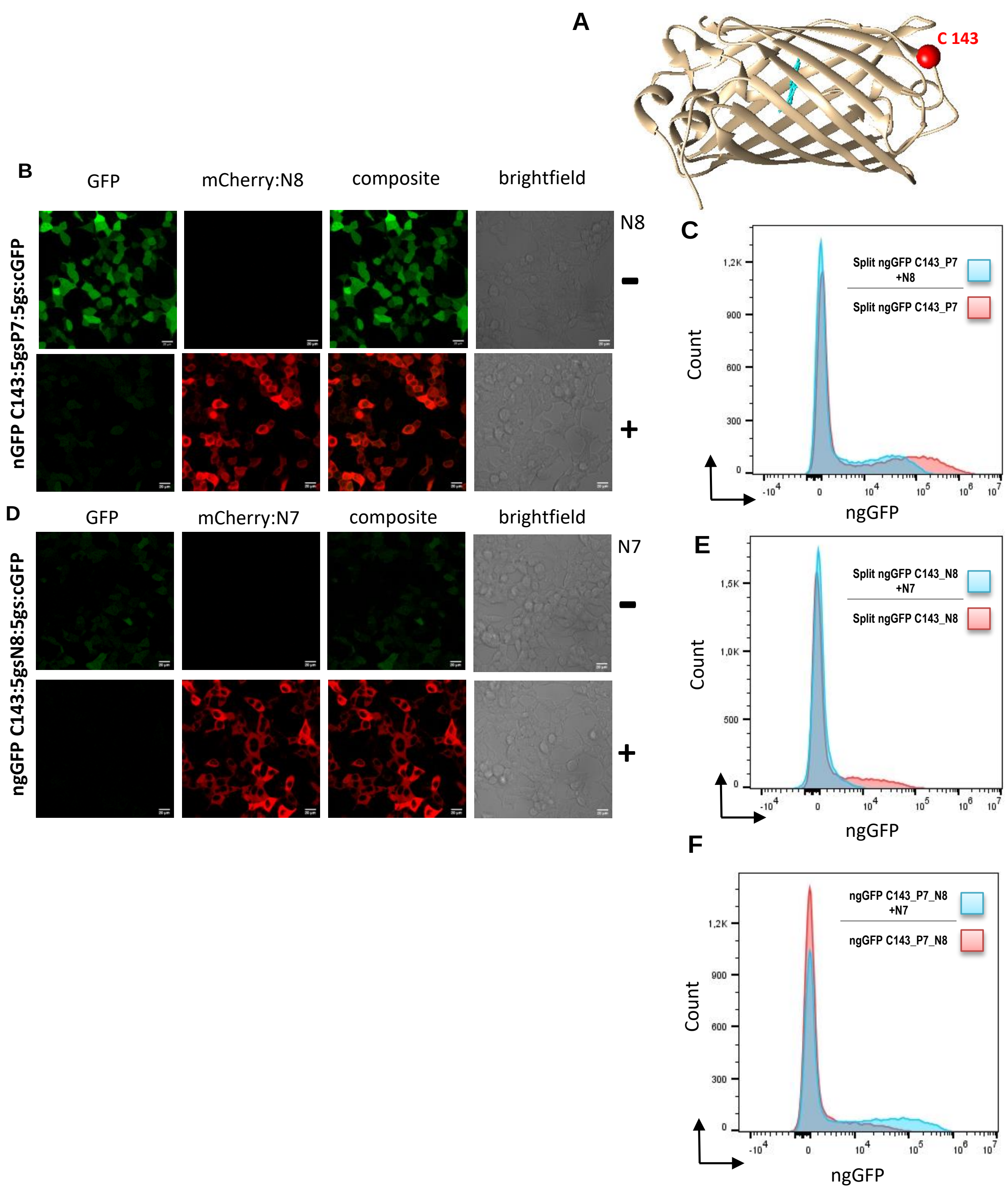


**Figure S5. Characterization of ngGFP INSRTR variant (OFF switch) with confocal microscopy. Related to Figure 5.**

**(A)** Structure of ngGFP with highlighted site of peptide insertion (red).

**(B)** ngGFP (50ng) with inserted peptide P7 at position C143; addition of mCherry:N8 (250 ng) deactivates ngGFP. **(C)** Flow cytometry analysis of fluorescent protein GFP (400ng) with inserted peptide P7 at position C143; addition of N8 (1000ng) deactivates ngGFP.

**(D)**ngGFP (50ng) with inserted peptide N8 at position C143; addition of mCherry:N7 (250ng) deactivates ngGFP.

Note: peptide N8 has higher helical propensity compared to P7. Insertion of N8 at position C143 thus reduces initial GFP fluorescence.

**(E)** Flow cytometry analysis of fluorescent protein GFP (400ng) with inserted peptide N8 at position C143; addition of N7 (1000ng) deactivates ngGFP.

**(F)** Flow cytometry analysis of fluorescent protein GFP (400ng) with inserted peptide P7 at position C143 and an autoinhibition loop N8; addition of N7 (1000ng) activates ngGFP. Experiments were made on transiently transfected HEK293T cells.

**RESOURCE AVAILABILITY**

**Lead Contact**

**Materials Availability**

Reagents generated in this study are available from the Lead Contact with a completed Materials Transfer Agreement.

**Data and Code Availability**

### Materials and Methods

**Molecular dynamics simulations (MDS)**

To gain insight into the structural dynamics of INSRTR structures, we preformed MDS with luciferase protein model using CHARMM27 all-atom force field embedded in GROMACS simulation package (*34*).

INSRTR models were prepared by inserting a loop into wildtype protein fold (PDB ID: 1BA3) using MODELLER (*35*). Protein models with coiled-coil structure were generated by extending the loop using GROMACS pull code. Umbrella pulling was applied between two residues representing the beginning and the end of a helix. Upon reaching the distance of 4 nm, coiled-coil structure was modelled on the loop backbone. For all the structures – solvated and electroneutral system was assembled, followed by energy minimization, equilibration under NVT (Constant number of particles, pressure and temperature) and NPT (Constant number of particles, volume and temperature) conditions. Equilibrated structures of luciferase were simulated for 5 ns and final MD trajectories were analyzed using GROMACS and Chimera (Pettersen et al., 2004)[36][36][36][36][36][36][36](*36*).

**AlphaFold2 modeling**

Protein models were built using publicly available scripts and modeling algorithm AlphaFold2 (*37, 38*)(Jumper et al., 2021; Mirdita et al.)[37, 38][37, 38][37, 38][37, 38][37, 38][37, 38][37, 38]. If AlphaFold2 was not able to generate a structure due its size or the structure had wrong pairing of CC, we prepared the models using MODELLER (*35*)(Eswar et al., 2007)[35][35][35][35][35][35][35].

**Plasmids and cell lines.**

All plasmids (listed in **Table S1**) were constructed using the Gibson assembly method. The human embryonic kidney cell line, HEK293T (ATCC CRL-3216), was cultured in complete media (DMEM; 1 g/l glucose, 2 mM l-glutamine, 10% heat-inactivated FBS (Gibco)) with 5% $CO_2$ at 37 ºC. We used plasmid pcDNA3 (Invitrogen) to express INSRTR proteins. *Renilla* luciferase (phRLTK, Promega) was used as a transfection control in the dual luciferase assay. Jurkat cell line was cultured in complete media (RPMI 1640 W/GLUTAMAX-I, Gibco, 10% heat-inactivated FBS (Gibco)) with 5% $CO_2$ at 37 ºC.

**INSRTR protein regulation.**

HEK293T cells were seeded in 96-well plates (Corning) at 2.5 × $10^4$ cells per well (0.1 ml). The next day, cells were transiently transfected with plasmids expressing split proteins (sequences are shown in Table S1), and phRL-TK (Promega) constitutively expressing *Renilla* luciferase (5 ng per well, to normalize transfection efficiency) using the PEI transfection reagent. The total amount of DNA for each transfection was kept constant by adding appropriate amounts of the control plasmid pcDNA3 (Invitrogen). For CC- regulated protein activity, HEK293T cells were transfected with plasmids expressing mCherry with C'- terminally fused coiled coil peptide under cytomegalovirus (CMV), and phRL-TK (Promega) constitutively expressing *Renilla* luciferase (5 ng per well) using PEI transfection reagent. The synthetic peptides were transfected into cells using DOTAP transfection reagent according to the manufacturer's instructions. Briefly, 24 h after plasmid transfection, peptides were diluted in Hepes buffer to the working concentration of 200 μM and then further diluted to final concentrations used on cells. The DOTAP transfection reagent was added to the peptide solution and transferred to the medium after 15-min incubation. Six hours later, the media was removed, and the cells were lysed using 1 × Passive lysis buffer.
The kinetics curves (Fig 2e, 3d) were fit to Velocity as a function of substrate (Nonlinear regression; allosteric sigmoidal) using GraphPad Prism 8 (Y=Vmax*X^h/(Khalf^h+ X^h)

Jurkat cells were electroporated using the Neon Electroporation System (Life Technologies). Prior to electroporation, cells were washed with PBS and resuspended in R buffer at a density of 3x$10^7$ cells per ml. 100 μL cells were mixed with 10 μg of plasmid DNA encoding the INSRTR variant of CD19 CAR and electroporated (1600 V, 3 pulses, 10 ms). Immediately after pulsing, cells were transferred to one well of a 12-well plate containing pre-warmed complete RPMI 1640 medium. The next day, 24h post electroporation, cells were counted and seeded in 96 well plate with target Raji cells at a ratio of Effector (Jurkat-CAR-T):Target (Raji)=5:1 and increasing amounts of a protein with inhibitory CC segment were added. Stimulation with target cells was terminated after 24hours by removal of media. Media was used for ELISA detection of interleukin-2 (hIL-2), produced as a result of Jurkat CAR-T cell activation.

**Flow cytometry**

Flow cytometry was used to determine expression levels of INSRTR modified CAR T variants on Jurkat cells. Cells were washed two times in buffer (2% BSA in PBS) before cell surface staining with fluorescent-labeled-antibody (anti-c-Myc 9B11 Alexa Flour 647 or anti-c-Myc 9B11 Alexa Flour 488). Cells were resuspended in 100 μL buffer containing antibody diluted 1:200 and incubated at 4°C for 30 min. Cells were washed twice with PBS and analyzed on a Cytek Aurora running SpectroFlo software. Flow cytometry data were analyzed with FlowJo v 10.8.1 software.

**Confocal microscopy**

For confocal microscopy, HEK-T 293 were transiently transfected with plasmids encoding INSRTR GFP and/or mCherry:CC. 48h after transfection, cells were analyzed. Microscopic images were acquired using the Leica TCS SP5 inverted laser-scanning microscope on a Leica DMI 6000 CS module equipped with an HCX Plane-Apochromat lambda blue 63 × oil-immersion objectives with NA 1.4 (Leica Microsystems, Wetzlar, Germany). A 488-nm laser line of a 100-mW argon laser with 10% laser power was used for GFP excitation, and the emitted light was detected between 500 and 540 nm. A 1-mW 543-nm HeNe laser was used for mCherry:CC excitation and emitted light was detected between 580 and 620 nm. The images were processed with LAS AF software (Leica Microsystems) and ImageJ software (National Institute of Mental Health, Bethesda, Maryland, USA).

**Dual luciferase assays.**

At indicated time points, the cells were lysed in Passive Lysis 1 × Buffer (Promega) and analyzed with a dual-luciferase reporter assay to determine the firefly luciferase and the *Renilla* luciferase activities

(Orion II microplate reader, Berthold Technologies). Relative luciferase units (RLU) were calculated by normalizing the firefly luciferase value to the constitutive *Renilla* luciferase value in each sample. Normalized RLU values (nRLU) were calculated by normalizing the RLU values of each sample to the value of the indicated sample within the same experiment.

**ELISA**

ELISA test was performed to determine secreted hIL2 from electroporated and stimulated Jurkat cells. High binding half-well plates (Greiner) were used. Human IL-2 was measured using standard ELISA assay according to manufacturer's protocol (hIL-2 ELISA Invitrogen 88-7025-88). In brief, plates were coated with primary antibody and incubated overnight (4 °C) Next day plates were washed with PBS+0,05 % Tween20 using ELISA plate washer (Tecan). Next, plates were blocked for 1h at RT with ELISA diluent (PBS+3%FBS) solution. Afterwards, plates were again washed. Then serial dilution of hIL2 standard and 1:2 diluted samples were added and incubated at RT for 2h. Next, plates were washed and afterwards detection antibody was added. Plates were incubated 1h at RT. Next plates were washed and HRP conjugated avidin was added and incubated for 30 min. After the addition of substrate (TMB solution) the reaction was stopped with 0.16 m sulfuric acid. The plates were read on a microplate reader at 450 nm, and again at 630 nm for correction by subtraction of the reading at 630 nm from that at 450 nm.

**Beta galactosidase assay.**

HEK293T cells transiently transfected with plasmids expressing β-galactosidase variants and N7 peptide using the PEI transfection reagent. Enzymatic activity of β-galactosidase was assessed with β-Gal Reporter Gene Assay, chemiluminescent (Roche, cat. no. 11758241001) according to manufactures' instructions. Briefly, 24h after transfection media was removed and cells were lysed with Lysis reagent. After 30min incubation, Substrate reagent was added for additional incubation of 30 min in the dark. Initiation of chemiluminescent reaction was achieved with automatic injection of Initiation solution with Orion II microplate reader (Berthold Technologies) following by luminescence readout. Enzymatic activities of β-galactosidase variants were plotted against standard curve of recombinant β-galactosidase from E. coli.

**Protein purification**

*Escherichia coli* NiCo21(DE3) strain (New England Biolabs) was transformed with constructs cloned in pET41a expression vector and grown at 37 °C overnight (160 r.p.m.) in Luria-Bertani (LB) medium containing kanamycin (50 μg/mL). Bacterial cultures were transferred to LB medium at OD of 0.1, grown at 37 °C until OD reached 0.6 and induced with 0.5 mM IPTG. After induction, the cultures were cultured for 4 additional hours at 30 °C. Cells were then harvested by centrifugation, pellets were frozen at -20 C overnight. Harvested cells were resuspended and lysed on ice with a lysis buffer: 50 mM Tris buffer at pH 8, 150 mM NaCl, 0.5 mg/mL Lysozyme (Sigma-Aldrich), 15 U/mL Benzonase (Millipore) and CPI protease inhibitor mix (Sigma-Aldrich). Cell lysis was completed by ultrasonication on ice for 6 min, at intervals of 1 s pulse and 3 s pause (45% amplitude). Subsequently, cellular lysates were centrifuged at 16,000g (4 °C) for 20 min, respective soluble fractions were filtered through 0.2 μm filter units (Sartorius) and applied to Ni-NTA resin (Golden Biotechnology) previously equilibrated with buffer (50 mM Tris buffer at pH 8.0, 150 mM NaCl, 10 mM Imidazole). After washing with buffer A and B (50 mM Tris buffer at pH 8.0, 150 mM NaCl, 20 mM imidazole), the bound fraction was eluted with buffer C (50 mM Tris buffer at pH 8.0, 150 mM NaCl, 250 mM imidazole). Glycerol (10% v/v) was added to the eluted fractions. The samples were then concentrated (Millipore centrifugal unit 3,5 MWCO), characterized by SDS-PAGE and shock-frozen in liquid nitrogen and stored at −80 °C.

**Statistical analysis**

All statistical analyses were performed using GraphPad Prism 8. All experiments were independently performed in triplicate unless otherwise indicated. Detailed statistic is listed in Table S2. All experiments showing representative data were repeated at least twice with similar results. Independent experiments refer to independent cell samples seeded, transfected, treated and analyzed on different days.

**SUPPLEMENTAL EXCEL TABLE TITLES AN LEGENDS**

**Table S1. Amino acid sequences of constructs used in this study. Related to methods.**

**Table S2. Detailed statistical analysis. Related to Figure 1-5.**
